# Disruption of VGLUT1 in cholinergic medial habenula projections increases nicotine self-administration

**DOI:** 10.1101/2021.06.19.449108

**Authors:** Elizabeth A. Souter, Yen-Chu Chen, Vivien Zell, Valeria Lallai, Thomas Steinkellner, William S. Conrad, William Wisden, Kenneth D. Harris, Christie D. Fowler, Thomas S. Hnasko

## Abstract

Cholinergic projections from the medial habenula (MHb) to the interpeduncular nucleus (IPN) have been studied for their complex contributions to nicotine addiction and have been implicated in nicotine reinforcement, aversion, and withdrawal. While it has been established that MHb cholinergic projections co-release glutamate, no direct evidence has demonstrated a role for this specific glutamate projection in nicotine consumption. In the present study, a novel floxed *Slc17a7* (VGLUT1) mouse was generated and used to create conditional knockout (cKO) mice that lack VGLUT1 in MHb cholinergic neurons. Histochemical approaches and optogenetics-assisted electrophysiology were used to validate the disruption of VGLUT1 from cholinergic MHb to IPN projections. The mice displayed no gross phenotypic abnormalities and exhibited normal exploratory and locomotor behavior in the open-field assay. However, the loss of VGLUT1-mediated glutamate co-release led to increased nicotine self-administration. These findings indicate that glutamate co-release from ventral MHb cholinergic neurons opposes nicotine consumption and provide additional support for targeting this synapse to develop potential treatments to nicotine addiction.

## INTRODUCTION

Despite decades of research demonstrating the negative consequences of smoking and emerging evidence on harmful effects of electronic cigarettes, the use of tobacco products persists. In the United States, nicotine use among adolescents has increased in recent years. For instance, in 2019, 23% of middle and high school students reported use of a nicotine-containing product in the past 30 days, up from 9% in 2014 [1,2]. As the main psychoactive and addictive compound, nicotine remains at the forefront of this continuing public health crisis [3-6].

Nicotine mediates its psychoactive effect by acting on nicotinic acetylcholine receptors (nAChRs) in the brain. Within the mesolimbic circuit, including ventral tegmental area dopaminergic projections to nucleus accumbens, activation of nAChRs contribute to the rewarding effect of nicotine [7-14]. Conversely, the excitatory projection from medial habenula (MHb) to interpeduncular nucleus (IPN) has been identified as a key substrate upon which nicotine actions contribute to nicotine aversion [15-24].

Both MHb and IPN express high levels of several nAChR subunits, which modulate excitability and neurotransmission within this projection [25-30]. Mice lacking α5-containing nAChRs self-administered significantly more nicotine at high (typically aversive) doses, an effect which was rescued by re-expression of α5 nAChR subunit to the MHb-IPN pathway, suggesting that α5-containing nAChRs here are necessary for nicotine aversion [19]. Additionally, overexpression of ?4 nAChR subunit leads to enhanced MHb activity and a strong aversive response to nicotine, which can be abolished by disruption of the α5 nAChR subunit in MHb [21]. More recently, it has been shown that knockdown of the α3 nAChR subunit in either the MHb or IPN increases nicotine intake in rats [24]. These data together suggest that nAChRs containing the α5, α3, and ?4 subunits mediate aversive signaling in the MHb-IPN. Further, IPN projections to laterodorsal tegmentum (LDTg) are also strongly modulated by nicotine, and inhibiting this projection is sufficient to block nicotine aversion [31]. Together, these findings establish the importance of MHb-IPN projections in modulating intake of nicotine and encoding its aversive properties.

MHb-IPN projections are topographically organized, with ventral MHb cholinergic projections targeting central IPN and Substance P projections from dorsal MHb targeting lateral IPN [32,33]. The cholinergic ventral MHb expresses nAChRs and has been particularly implicated in nicotine aversion [19,21,26,28-30,34]. However, these neurons also express vesicular glutamate transporters (VGLUT) and can thus co-release both ACh and glutamate [20,35,36], raising the question of what the glutamate signal from MHb may contribute to nicotine intake. To address this question, we made a new conditional VGLUT1 mouse line and used it to generate conditional knock-out (cKO) mice that lack VGLUT1 in ventral MHb cholinergic neurons. We show that cKO mice have reduced glutamate transmission in MHb projections to IPN and that cKO mice display increased intravenous (IV) nicotine self-administration, consistent with a role for VGLUT1-mediated glutamate co-release at this circuit in opposing nicotine intake.

## METHODS

### Animals

Mice were used in accordance with the University of California, San Diego and the University of California, Irvine Institutional Animal Care and Use Committees. BAC transgenic Kiaa1107-Cre mice were obtained from GENSAT through the MMRRC (#034692-UCD). Kiaa-Cre mice were bred hemizygously with C57Bl/6J wildtype mice (Jackson Laboratory, 000664). VGLUT2-IRES-Cre (*Slc17a6*^*Cre*^) knock-in mice were ordered from Jackson Laboratory (#028863) and bred homozygously or to C57BL6/J wildtype mice. All experiments were done in adult mice (aged greater than 8wks) and in both males and females in approximately equal proportion.

### VGLUT1 conditional allele

To generate VGLUT1-floxed mice (*Slc17a7*^flox^), a targeting vector containing two loxP sites flanking *Slc17a7* exons 4-7 and an FRT-flanked neomycin (Neo) resistance cassette was electroporated into C57Bl/6-derived ES cells. Antibiotic (G418) resistant colonies were selected, isolated, and amplified. The amplified clones were screened for homologous recombination at the *Slc17a7* locus by PCR. Southern blot analysis was used to confirm both 3’ and 5’ homologous recombination. Blastocysts were isolated from pregnant C57Bl/6J-Tyr^c-2J^/J (albino C57Bl/6) females, injected with one of six validated ES cell clones, and implanted into pseudo-pregnant females. Chimeric males were bred to C57Bl/6 females with constitutive expression of FLP recombinase to excise the Neo cassette in F1 offspring. F1 mice were crossed to C57Bl/6, excision was confirmed by PCR and Southern blot, and these F2 mice were used to establish the VGLUT1 floxed line (*Slc17a7*^*flox*^). To generate conditional knock-out mice (*Kiaa*^*Cre*^; *Slc17a7*^*flox/flox*^), Kiaa^Cre^ mice were bred to homozygous *Slc17a7*^*flox/flox*^ and resulting heterozygotes (*Kiaa*^*Cre*^; *Slc17a7*^*+/flox*^) were then bred to homozygous *Slc17a7*^*flox/flox*^ mice. Mice were group housed and maintained on a 12-hour light-dark cycle. Food and water were available *ad libitum* except where noted.

### Stereotactic surgery

For intracranial injections, mice (>4 weeks) were deeply anesthetized with isoflurane, placed in a stereotaxic frame (Kopf), and bilaterally injected with AAV1-Ef1a-DIO-ChR2:mCherry (2 × 10^12, UNC Gene Therapy Center) into the MHb (LM = -1.15, AP = -1.58, DV = -2.42 and -2.00, 20° angle; right: LM = +0.95, AP = -1.58, DV = -2.42 and -2.00, 20° angle; mm relative to *Bregma*). Two 150nL aliquots were given per hemisphere at 100nl/min using pulled glass pipettes (Nanoject III, Drummond Scientific). Analgesic was given before and at least one day after surgery (Carprofen, Zoetis, 5mg/kg s.c.). Mice were monitored daily for 5d after surgery and allowed to recover for at least 21d before histological processing or 28d before electrophysiological recordings.

### Immunohistochemistry

Mice were deeply anesthetized with pentobarbital (200mg/kg s.c., VetOne) and transcardially perfused for 2m with ice-cold phosphate buffered saline (PBS) then for 8 min with ice-cold 4% paraformaldehyde (PFA) at a rate of 5-6ml/min. Brains were prepared as previously described [37]. Primary antibodies used: DsRed (Rabbit, 1:2000, Clontech), VGLUT1 (Guinea Pig, 1:2000, Synaptic Systems), VGLUT2 (Rabbit, 1:1000, Synaptic Systems), ChAT (Goat, 1:200, Millipore). Secondary antibodies used (5ug/ml, Jackson ImmunoResearch): Alexa488 Donkey Anti-Goat (705-545-147), Alexa 488 Donkey Anti-Guinea Pig (706-545-148), Alexa594 Donkey Anti-Guinea Pig (706-585-148), Alexa 594 Donkey Anti-Rabbit (711-585-152), Alexa647 Donkey Anti-Rabbit (711-605-152), Alexa647 Donkey Anti-Goat (705-605-147). Images were captured using a Zeiss AxioObserver Z1 epifluorescence microscope (10x 0.45NA, 20x 0.75NA, or 63x 1.4NA objective) and Zen software. Densitometry was done with Fiji/ImageJ.

### Fluorescent *in situ* hybridization

Mice were deeply anesthetized with pentobarbital before cervical dislocation. Brains were prepared as previously described [37]. In situ hybridization was done using RNAscope Multiplex Fluorescent Assay (Advanced Cell Diagnostics) according to manufacturer specification. *Slc17a7* (503511), *ChAT* (408731-C2), and *Cre* (312281-C3) were coupled to Atto550, Alexa647, and Alexa488, respectively and counterstained with DAPI. Images were captured using a Zeiss AxioObserver Z1 epifluorescence microscope and processed with Zen software.

### Open-Field Behavior

Mice were placed in an Open Field (30m) measuring 50×50 cm and their activity was recorded and analyzed using AnyMaze software (San Diego Instruments). The field was cleaned with 70% ethanol between sessions. The field was segmented into a 5×5 grid, with the innermost 9 squares designated as the center.

### Operant Behavior

Mice were mildly food-restricted to 85-90% of their free-feeding weight and were then trained to lever press for food pellets (grain-based, 20 mg, 5TUM, TestDiet) on a two-lever operant task across ascending fixed ratio (FR) schedules from 1 up to 5 lever presses, as previously described [18]. At the start of the session, both levers were extended into the chamber and were present throughout the 1-hr session. Responses on the active lever that met the FR criteria resulted in the delivery of a food pellet, which was paired with a cue light for a 20-s time-out period, resulting in the final reinforcement schedule of FR5TO20 for food training sessions 4-7. Responses on the inactive lever were recorded but had no scheduled consequences. Testing was conducted 6-7d/wk and behavioral responses were recorded with a MedPC interface (Med Associates). Thereafter, subjects were anesthetized (isoflurane) and catheterized as previously described [18,38]. The catheter tubing was passed subcutaneously from the animal’s back to the right jugular vein, a 1-cm length of catheter tip was inserted into the vein and tied with surgical silk suture. Following surgery animals were allowed ≥48 hr to recover, and were then provided 1-hr access to re-establish food responding under the FR5TO20 sec schedule until the criteria of >30 pellets/session were again achieved. Mice were then transitioned to respond for intravenous nicotine self-administration in lieu of food using the same FR5TO20 sec, 1-hr daily sessions, 6-7d/wk, at the training dose of nicotine (0.03 mg/kg/infusion) for 8 d. For all doses, nicotine (0.03 ml per infusion volume) was delivered through tubing into the intravenous catheter by a Razel syringe pump (Med Associates). After achieving stable responding on the 0.03 mg/kg/infusion dose, mice were transitioned to the moderate dose of 0.1 mg/kg/infusion nicotine for 5d. This dose results in a similar levels of drug intake as that found at higher doses with behavioral titration via self-administration and was used to further establish baseline responding in between access to each subsequent varying dose [18]. Next, the mice were provided access to either the low 0.01 mg/kg/infusion or high 0.4 mg/kg/infusion dose for 5d, and then re-established at baseline on 0.1 mg/kg/infusion for at least 2d, and thereafter given access to the counterbalanced doses of either 0.01 or 0.4 mg/kg/infusion for an additional 5d. Following re-establishing baseline for at least 2d, the mice were then provided access to respond for saline vehicle. The mean of the final 3 days on each dose was calculated for each subject. Catheters were flushed daily with physiological sterile saline solution (0.9% w/v) containing heparin (100 units/mL). Catheter integrity was verified with the ultra-short-acting barbiturate anesthetic Brevital (2%, methohexital sodium, Eli Lilly) at the end of the study.

### Electrophysiological recordings

Recordings were performed on adult mice (7-12wks) as previously described [39]. mCherry-labeled MHb terminals were visualized by epifluorescence and visually guided patch recordings were made using infrared-differential interference contrast (IR-DIC) illumination (Axiocam MRm, Examiner.A1, Zeiss). ChR2 was activated by flashing blue light (473 nm) through the light path of the microscope using a light-emitting diode (UHP-LED460, Prizmatix) under computer control. Neurons were held in voltage-clamp at -60 mV to record EPSCs in whole-cell configuration and single-pulse photostimuli (5-ms or 1-s pulse width) were applied every 45 s and 10 photo-evoked currents were averaged per neuron per condition. Stock solutions of DNQX (10 mM in DMSO, Sigma) and mecamylamine hydrochloride (10 mM, Tocris) were diluted 1,000-fold in ACSF and bath applied at 10 μM. Current sizes were calculated by using peak amplitude from baseline.

### Statistics

Data analysis was done using GraphPad Prism v9. Data were analyzed using t-test corrected for multiple comparisons (Bonferroni-Sidak)(figures 2b, 2c, 2d), unpaired t-test (4c, 4d, 5f), RM two-way ANOVA (Sidak post hoc)(5b, 5c, 5g), or RM three-way ANOVA (Sidak post hoc)(5d, 5e).

## RESULTS

### Generation of *Slc17a7* conditional knock-out from ventral MHb

To target cholinergic neurons of the MHb we used the Kiaa1107-Cre (*Kiaa*^*Cre*^) transgenic line that has been used previously to disrupt Choline acetyltransferase (ChAT) expression in MHb [20]. The functional expression of *Cre recombinase* driven by *Kiaa1107* regulatory elements was first validated by crossing to the Ai6 ZsGreen reporter [40] to generate *Kiaa*^*Cre*^; *Rosa26*^*ZsGreen*^ mice (**figure 1a**). Robust ZsGreen fluorescence was seen in MHb, with densest expression observed in ventral (basolateral and basomedial) MHb (**figure 1b**).

**Figure 1.**
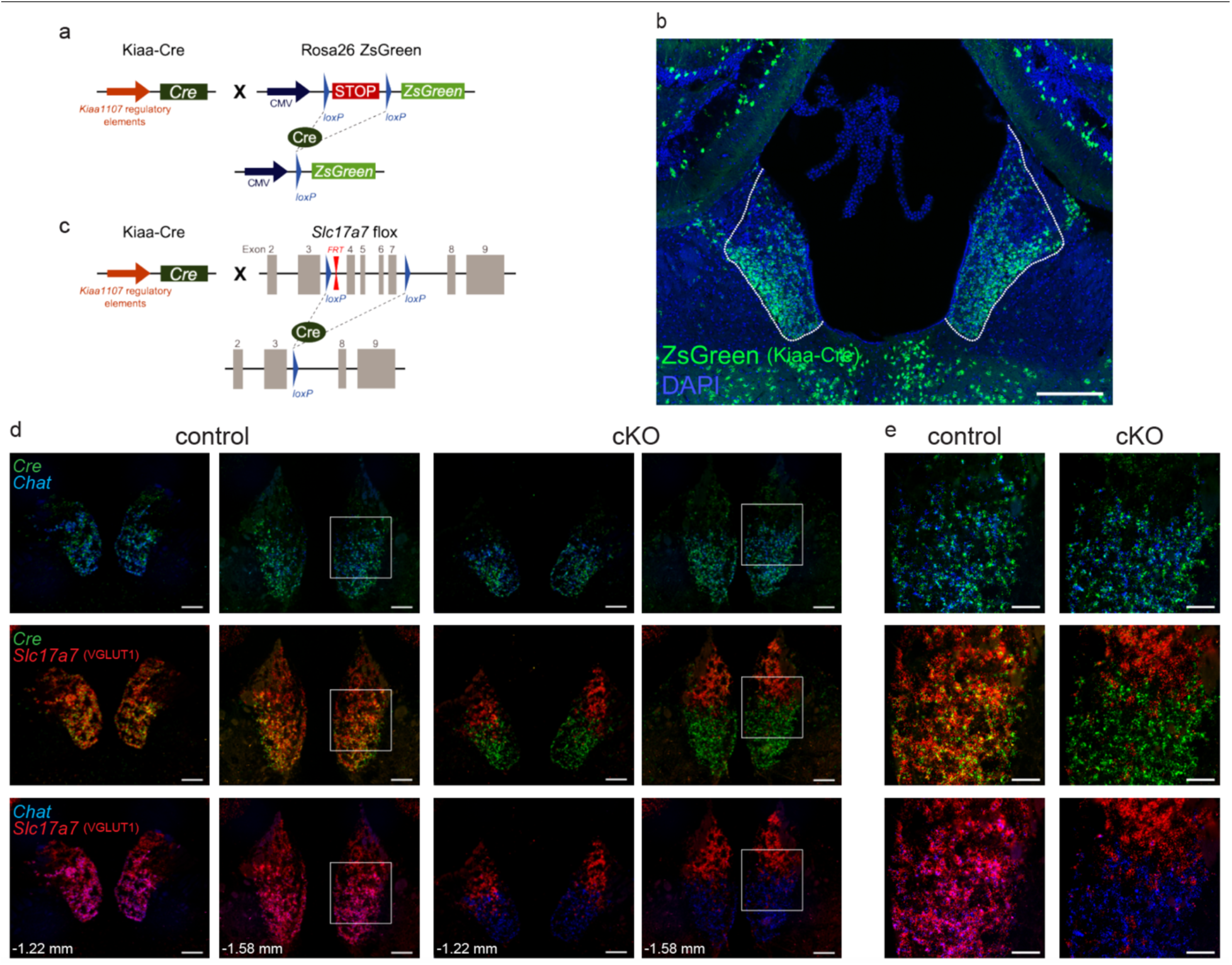
Conditional knock-out of *Slc17a7* (VGLUT1) from cholinergic neurons in MHb. **(a)** Schematic of *Kiaa*^*Cre*^; *Rosa26*^*ZSGreen*^ reporter mouse line with Cre expression driven by *Kiaa1107* regulatory elements and ZsGreen expression dependent on Cre recombination. **(b)** Native ZsGreen fluorescence counterstained with DAPI in MHb (outlined); scale 200 μm. **(c)** Schematic of Cre recombination of the floxed *Slc17a7* (VGLUT1) locus in the cKO (*Kiaa*^*Cre*^; *Slc17a7*^*flox/flox*^) mouse line. **(d)** Fluorescent *in situ* hybridization of *Cre, Chat*, and *Slc17a7* expression in MHb of control and cKO mice at two Bregma points; scale 100 μm. **(e)** Higher magnification images from white squares in (d); scale 50 μm.

**Figure 2:**
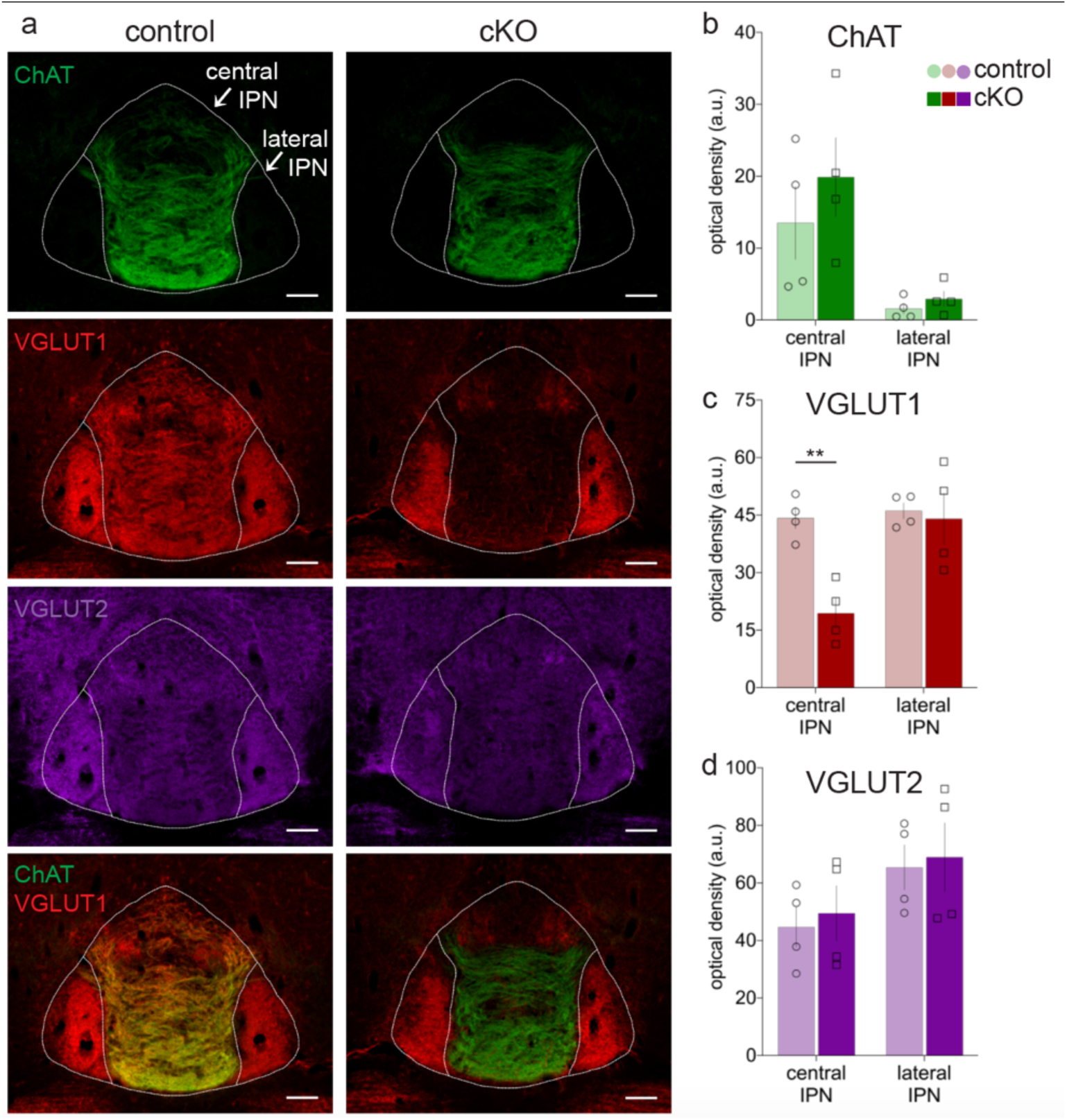
Loss of VGLUT1 in central IPN of cKO mice. **(a)** Immunohistochemistry for ChAT, VGLUT1, and VGLUT2 in the IPN of control (*Slc17a7*^*flox*^) and cKO mice (*Kiaa*^*Cre*^; *Slc17a7*^*flox/flox*^); bottom row shows ChAT and VGLUT1 merge; scale 100 μm. Densitometric quantification in central and lateral IPN of (**b**) ChAT, (**c**) VGLUT1, and (**d**) VGLUT2 signals. Only VGLUT1 was significantly reduced in cKO and only in the central IPN (*\*\*p=0*.*0040*); n=4 mice per group.

To disrupt VGLUT1 expression, we generated a novel mouse line carrying a VGLUT1 conditional allele (*Slc17a7*^*flox*^) with exons 4-7 flanked by loxP sites (**figure 1c**). We next crossed *Slc17a7*^*flox*^ mice to *Kiaa*^*Cre*^ to generate the VGLUT1 conditional knockout (cKO, *Kiaa*^*Cre*^; *Slc17a7*^*flox/flox*^). We then generated an RNAscope probe targeting exons 4-7 of *Slc17a7* and used this together with probes against a cholinergic marker (*Chat*) and *Cre recombinase* on sections from cKO and control (*Kiaa*^*Cre*^) mice (**figure 1d-e**). The pattern of *Cre* expression was identical for both genotypes and similar to the pattern observed in the ZsGreen reporter cross, with robust expression in ventral MHb. *Chat* appeared unchanged across genotype and showed high overlap with *Cre. C*onsistent with other reports, *Chat* expression was largely restricted to ventral MHb [35,41-43].

Also consistent with other previous reports, *Slc17a7* (VGLUT1) was expressed throughout the MHb in controls [35,44-46]. However, the cKO showed a markedly different pattern. In cKO mice, *Slc17a7* (VGLUT1) was virtually eliminated from ventral MHb, but expression was intact in dorsal (apical) MHb (**figure 1d-e**). These results indicate that our cKO successfully and selectively disrupted *Slc17a7* expression from Cre-expressing neurons in ventral MHb.

We next used immunohistochemistry to examine the expression of pre-synaptic cholinergic and glutamatergic markers in the IPN, the major projection target of MHb (**figure 2a**). While the expression of ChAT was unaffected in the cKO (**figure 2b**), the pattern of VGLUT1-labeled fibers was markedly different depending on genotype and subregion. cKO mice had a significant reduction of VGLUT1 expression compared to controls in central IPN (*t(6)=5*.*2, p(adj)=0*.*004*), but no difference in VGLUT1 between genotypes was observed in lateral IPN (*t(6)=0*.*30, p(adj)=0*.*95*) (**figure 2c**). Together, these data are concordant with our RNAscope data and demonstrate the selective disruption of VGLUT1 from cholinergic MHb inputs that target the central region of the IPN, which includes the caudal, dorsomedial, intermediate, and rostral subnuclei.

We also examined VGLUT2-labeled fibers in the IPN to test whether the loss of VGLUT1 led to changes in VGLUT2 expression. We detected no significant difference in VGLUT2 expression between genotypes in either central IPN (*t(6)=0*.*41, p(adj)=0*.*91*) or lateral IPN (*t(6)=0*.*25, p(adj)=0*.*96*) (**figure 2d**). These data argue against compensatory change in VGLUT2 expression following loss of VGLUT1 from cholinergic neurons in MHb.

### Expression of *Slc17a6* (VGLUT2) in IPN-projecting MHb neurons

The absence of *Slc17a7*/VGLUT1 expression in ChAT-expressing neurons of ventral MHb and central IPN provides strong evidence for loss of VGLUT1-mediated glutamatergic co-release from cholinergic MHb inputs in cKO mice. But while VGLUT1 has been implicated in mediating glutamate co-release from MHb cholinergic neurons [20,35,36], there is also evidence that some MHb neurons express *Slc17a6*/VGLUT2 [35,45,46], consistent with the presence of VGLUT2-labeled fibers that we observed in IPN (**figure 2d**). To directly test whether *Kiaa*^*Cre*^-expressing MHb cells also express *Slc17a6* (VGLUT2) we used RNAscope. *Slc17a6* was observed throughout the MHb (**figure 3a**), including in the ventral MHb where it partially co-localized with Cre recombinase (*Kiaa*^*Cre*^) (**figure 3b**).

**Figure 3:**
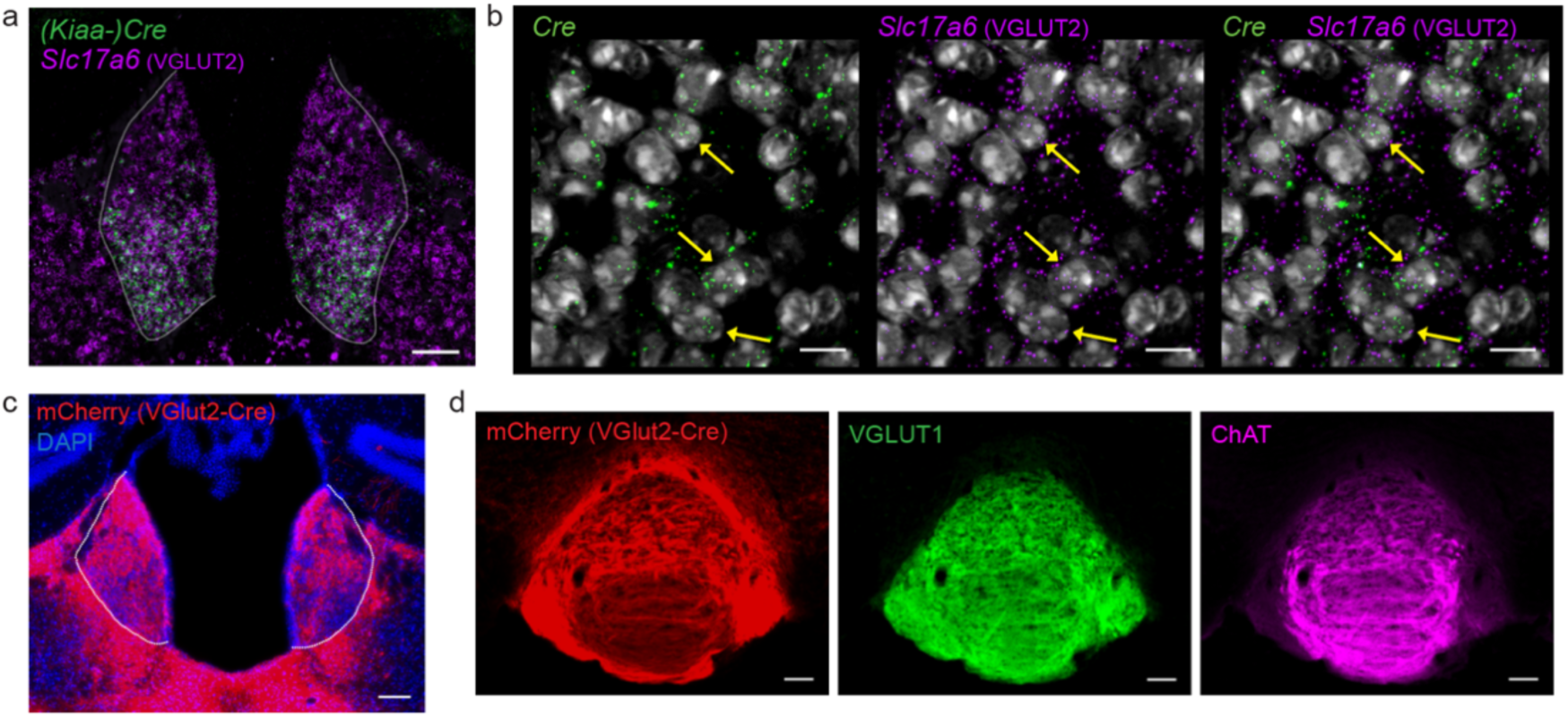
VGLUT2-expressing projections from MHb to IPN. **(a)** Fluorescent *in situ* hybridization from *Kiaa*^*Cre*^ mouse showing *Cre* and *Slc17a6* expression in the MHb (outlined); scale 100 μm. **(b)** High-resolution image showing expression of *Cre, Slc17a6* (VGLUT2), and DAPI in MHb of *Kiaa*^*Cre*^ mouse; scale 10 μm. Yellow arrows indicate some of the cells containing both *Cre* and *Slc17a6* mRNA. **(c)** Image of MHb from *Slc17a6*^*Cre*^ (VGLUT2) mouse injected with AAV1-Ef1α-DIO-ChR2:mCherry bilaterally into the MHb (outlined); scale 100 μm. **(d)** Immunohistochemistry of IPN from *Slc17a6*^*Cre*^ (VGLUT2) mouse injected bilaterally with AAV1-Ef1α-DIO-ChR2:mCherry in MHb. VGLUT2^Cre^ MHb terminals in IPN represented by mCherry+ expression, stained with VGLUT1 and ChAT; scale 100 μm.

The presence of MHb neurons co-expressing *Cre* and *Slc17a6* (VGLUT2) in cKO mice raised the question of whether this VGLUT2 population projected to IPN. We thus injected an Adeno-associated virus (AAV) into the MHb of *Slc17a6*^*Cre*^ (VGLUT2) mice [47] to Cre-dependently express Channelrhodpsin-2 fused to a fluorescent tag (ChR2:mCherry). Three weeks after surgery mCherry expression was found in MHb, as well as in surrounding areas of lateral habenula (LHb) and paraventricular nucleus of the thalamus (PV) (**figure 3c**). mCherry expressing fibers, presumably axon terminals from MHb, were also present in both central and lateral IPN (**figure 3d**). These results indicate that at least some Kiaa^Cre^ cholinergic neurons in MHb could express both VGLUT1 and VGLUT2, consistent with previous reports of VGLUT1/VGLUT2 co-expression in MHb [20,35].

### Disruption of VGLUT1 from ventral MHb neurons decreased evoked glutamate currents in central IPN

We next tested how the loss of VGLUT1 from Cre-expressing ventral MHb cholinergic neurons affected glutamate transmission from terminals in central IPN. We expressed ChR2:mCherry in MHb as above, but now using *Kiaa*^*Cre*^ and cKO mice that lack VGLUT1 in these neurons (**figure 4a**). ChR2:mCherry expression was observed in MHb and IPN (**figure 4b**); recordings were made from IPN neurons, with optogenetic stimulation of MHb terminals. Whole-cell voltage-clamp was used to assess optogenetic evoked excitatory postsynaptic currents (oEPSC) in response to either a single pulse of blue light (5ms) or train stimulation (5ms pulses at 20Hz for 1s). Single-pulse stimulation evoked DNQX-sensitive fast glutamatergic oEPSCs (**figure 4c**); oEPSCs were significantly smaller but not eliminated in the cKO (*unpaired t-test; t(14)=4*.*0, p=0*.*001*). Residual currents were presumably due to expression of VGLUT2 in some of the Cre-expressing cholinergic neurons and were blocked by bath application of an AMPA-type glutamate receptor antagonist. Train stimulation led to a mixed response that contained both faster glutamatergic, as well as slower cholinergic oEPSCs (**figure 4d**) that were blocked by a nicotinic acetylcholine receptor antagonist. While the variability in responses to train stimulation appeared higher in the cKO, no significant difference in oEPSC amplitude was detected between genotypes in response to train stimulation (*unpaired t-test; t(13)=1*.*6, p=0*.*14)*.

**Figure 4:**
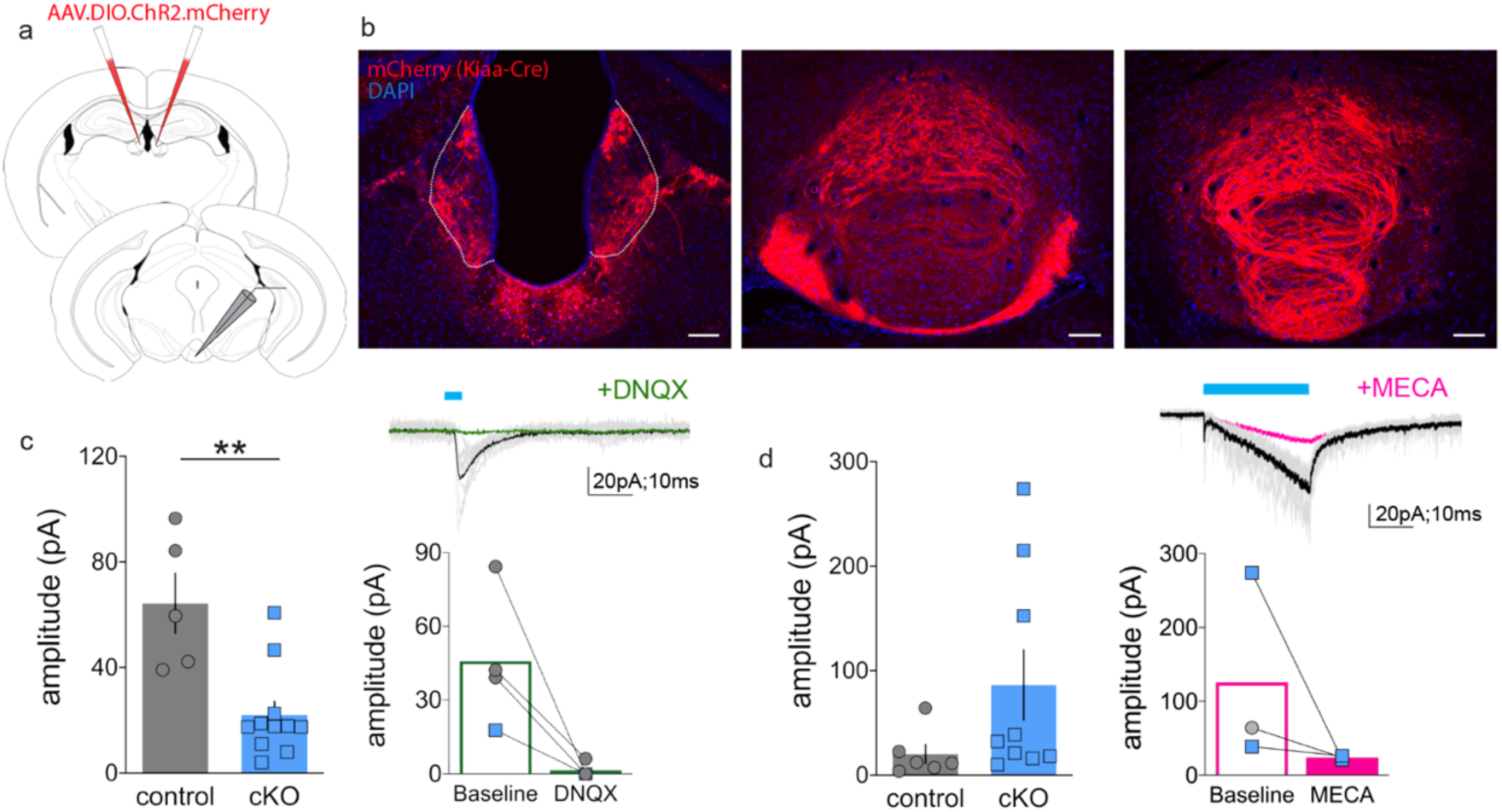
Reduced glutamate transmission from MHb to IPN in VGLUT1 cKO mice. **(a)** Schematic of electrophysiological preparation, with bilateral injections of AAV1-Ef1α-DIO-ChR2:mCherry in MHb of control (*Kiaa*^*Cre*^) or cKO (*Kiaa*^*Cre*^; *Slc17a7*^*flox/flox*^) mice. Slice recordings using optogenetic stimulation performed in the IPN 3+weeks after injection. **(b)** Images from control mouse of native Cre-dependent mCherry fluorescence in MHb (left) and fibers in IPN (center, right); scale 100 μm. **(c)** Whole-cell recordings in IPN with single-pulse optogenetic stimulation of MHb terminals led to oEPSC amplitudes that were reduced in the cKO (left, \*\**p=0*.*001*); blocked by bath application of DNQX (right). **(d)** oEPSC amplitude following train stimulation (1s) did not differ significantly different between control and cKO groups (*left*); sensitive to mecamylamine (right).

### Loss of MHb VGLUT1 increased nicotine self-administration

Prior studies have implicated MHb cholinergic signaling to IPN in the aversive effects of nicotine [17,18,20-22,34], but the contribution of glutamate co-release from this circuit had not been examined. We thus assessed the behavioral phenotype of littermate control and cKO mice. To test gross locomotor and exploratory behavior, we assessed mice in the open-field test. No significant differences were found between genotype in distance traveled (*RM two-way ANOVA; main effect of segment, F(2*.*6,36)=30, p<0*.*0001; genotype, F(1,14)=0*.*12, p=0*.*73; segment x genotype interaction, F(5,70)=2*.*0, p=0*.*084*) (**figure 5a**) or time spent in center (*main effect of segment, F(3*.*4,47)=1*.*1, p=0*.*38; genotype, F(1,14)=0*.*14, p=0*.*71; segment x genotype interaction, F(5,70)=0*.*19, p=0*.*97*) (**figure 5b**).

**Figure 5:**
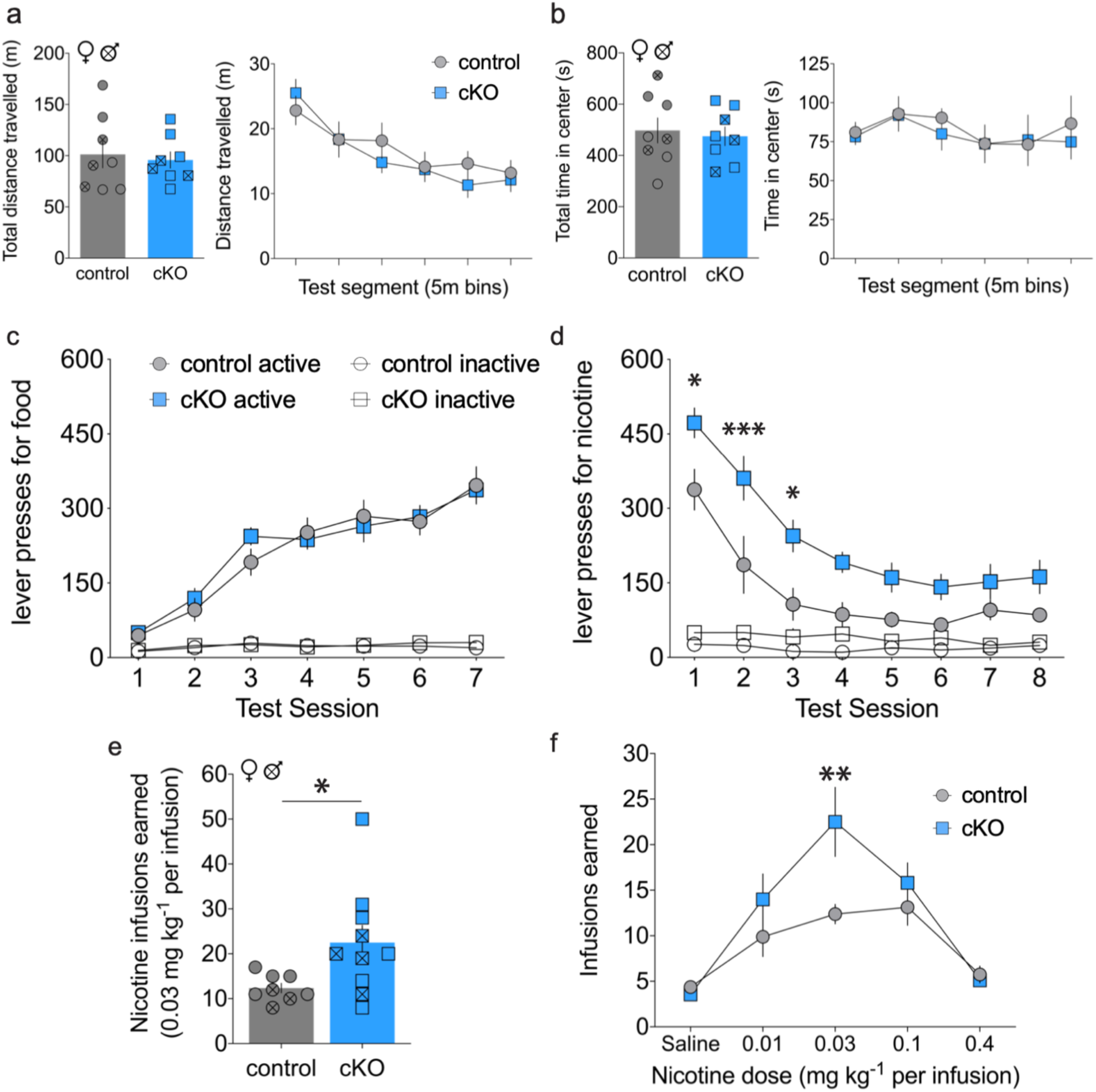
Increased nicotine self-administration in cKO mice. **(a)** Total distance traveled (*left*) and distance travelled across test segments (*right*) in open field assay showed no significant differences between genotype. **(b)** Total time in center (*left*) and time in center across test segments (*right*) did not differ between genotype. **(c)** Active and inactive lever presses during food training across test session did not differ by genotype. **(d)** Active and inactive lever presses for nicotine show increased self-administration for cKO mice (*Sidak’s *p<0*.*05, ***p<0*.*001)*. **(e)** cKO mice earned more total nicotine infusions in first 3 nicotine test sessions (t-test, *\*p<0*.*05)*. **(f)** Nicotine infusions earned by control and cKO mice in dose-response paradigm (*Sidak’s, **p<0*.*01)*.

Next, control and cKO mice were trained to lever press for food pellets and each reward delivery was paired with a cue-light for a 20-s timeout period (TO20). Across the initial 3 sessions, the fixed ratio (FR) schedule increased from 1 to 5 lever presses, then mice were maintained on an FR5 for an additional 3 sessions. No significant differences between genotypes were detected, suggesting intact operant learning and lever discrimination in cKO mice (*RM three-way ANOVA; main effect of session, F(6,108)=72, p<0*.*0001; genotype, F(1,18)=0*.*10, p=0*.*75; lever, F(1,18)=243, p<0*.*0001; session x genotype, F(6,108)=0*.*90, p=0*.*50; session x lever, F(6,108)=74, p<0*.*0001; genotype x lever, F(1,18)=0*.*026, p=0*.*87; session x genotype x lever, F(6,108)=1*.*3, p=0*.*25*) (**figure 5c**).

After food training, intravenous catheters were implanted, and an acquisition dose of nicotine (0.03 mg/kg/infusion) was introduced at the established FR5 TO20 schedule of reinforcement. As previously observed with this protocol [18], both groups pressed initially at a high rate similar to that observed with food reinforcement, which subsequently declined across sessions to a steady-state rate of nicotine self-administration (**figure 5d**). Interestingly, cKO mice engaged in consistently higher levels of nicotine self-administration across sessions at this dose *(RM three-way ANOVA; main effect of session, F(7,112)=38, p<0*.*0001; genotype, F(1,16)=11, p=0*.*004; lever, F(1,16)=111, p<0*.*0001; session x genotype, F(7,112)=1*.*8, p=0*.*099; session x lever, F(7,112)=33, p<0*.*0001; genotype x lever, F(1,16)=8*.*5, p=0*.*010; session x genotype x lever, F(7,112)=0*.*95, p=0*.*47*). Compared to controls, cKO mice earned significantly more total nicotine infusions in the first three test sessions *(unpaired t-test; t(16)=2*.*3, p=0*.*035*) (**figure 5e**).

To assess across a range of nicotine doses, a dose-response was then performed. While both groups exhibited an inverted U-shaped dose-response, cKO mice showed a dose-dependent increase in nicotine consumption compared to controls and this effect was most pronounced at 0.03 mg/kg/infusion dose *(RM two-way ANOVA; main effect of dose, F(4,64)=24, p<0*.*0001; genotype, F(1,16)=2*.*4, p=0*.*14; dose x genotype interaction, F(4,64)=3*.*5, p=0*.*012)* (**figure 5f**). Together, these results support the hypothesis that VGLUT1-mediated glutamate transmission from MHb to IPN opposes nicotine self-administration.

## DISCUSSION

Nicotine has both positively and negatively reinforcing properties. The balance of these two vary by administration route, dose, prior nicotine exposures, as well as by context, cues, schedules and contingencies [48-51]. Negative somatic and psychological effects of nicotine include nausea, dizziness, and anxiety [52,53]. Humans and monkeys will work to delay non-contingent intravenous infusions of nicotine, and higher doses can elicit conditioned place aversion in mice [54,55].

MHb projections to IPN are critical regulators of nicotine consumption. Nicotine facilitates glutamate release from MHb terminals by activating pre-synaptic α5-containing nAChRs [25,56]. Selective knock-down of α5 nAChR in MHb led to increased nicotine consumption in mice, as did blocking glutamate transmission in IPN by microinjection of NMDA-receptor antagonist [19]. More recently, targeted knock-down of α3 nAChR subunit in either MHb or IPN was shown to produce similar increases in nicotine intake [24]. Global overexpression of β4 nAChR subunit led to increased nicotine aversion, an effect reversed by selective expression of α5 nAChR subunit in MHb [21]. Together these data indicate that nicotine acting on α5-, α3-, and ?4-containing nAChRs facilitates nicotine-mediated excitatory transmission at MHb synapses in the IPN, which reduces nicotine-self administration. Importantly, our data demonstrate that cKO of VGLUT1 in the MHb led to increased nicotine self-administration, which is consistent with this framework and provide the first direct evidence that release of glutamate from cholinergic MHb projections to IPN inhibits nicotine consumption.

The MHb is a heterogenous structure composed of several unique cell types, each capable of releasing or co-releasing a variety of neurotransmitters and neuropeptides [57,58]. For example, Fos data indicate that most MHb cell types are activated by foot-shock stress [57]. On the other hand, activity in dorsal MHb neurons, which are largely non-cholinergic, may play a role in positive reinforcement and reward consumption [59,60]. Further, stimulation of glutamatergic septal inputs to MHb was anxiolytic, though different populations of MHb neurons were either inhibited or excited by this manipulation [61]. Thus, different MHb cell types appear to play opposing roles in mediating behaviors and affective states relevant to nicotine consumption.

Disruption of glutamate transmission from MHb to IPN could increase nicotine self-administration if this glutamate signal opposes nicotine reward or mediates aspects of nicotine aversion, though several lines of evidence favor the latter. For example, MHb projections to the IPN mediate negative affective behaviors such as anxiety, aversion, and the expression and extinction of fear memories [19,31,61-65]. In mice undergoing nicotine withdrawal, optogenetic silencing of MHb inputs to IPN reduced marble-burying and increased time spent in the open arms of an elevated plus maze; microinjection of NMDA antagonist in IPN recapitulated this effect and was also shown to reduce somatic signs of withdrawal [66,67]. Therefore, the loss of glutamate release from cholinergic MHb projections in our study most likely led to increased nicotine consumption by reducing its aversive properties.

Previous reports have detailed activation of the central IPN by MHb projections via fast glutamate-mediated currents, as well as by slower ACh-mediated currents [20,25,36]. Histological assessments of VGLUT1 and the vesicular ACh transporter (VAChT) have shown MHb axon terminals co-positive for these transporters, and electron microscopy has identified vesicles at this synapse containing both vesicular transporters [20,35,36]. Our experiments show presence of both *Slc17a7*/VGLUT1 and *Slc17a6*/VGLUT2 RNA transcripts in MHb, and that VGLUT2-expressing MHb neurons can also project to both lateral and central IPN, consistent with prior findings [57,58,68]. In our VGLUT1 cKO animals, the residual glutamate-mediated oEPSCs still present in IPN are most likely facilitated by expression of VGLUT2. Nevertheless, cKO of VGLUT1 led to a large reduction in evoked glutamate currents and to decreased nicotine intake, though disrupting both vesicular glutamate transporters might produce an even larger effect which may be addressed in future studies.

Work by Frahm and colleagues used a similar approach to disrupt ChAT expression in MHb and showed that this led to loss of nicotine withdrawal behaviors and loss of nicotine conditioned place preference [20]. Thus, despite both transmitters exerting primarily excitatory post-synaptic actions, glutamate and ACh release from MHb neurons appear to mediate different affective responses to nicotine – with cholinergic transmission necessary for nicotine-associated reward, and glutamate transmission signaling nicotine aversion. This is perhaps more surprising given that these transmitters localize to an overlapping pool of synaptic vesicles and synergistic effects on vesicle filling are supported by data demonstrating that loss of ChAT/ACh reduces glutamate filling [20], presumably because ACh uptake through VAChT dissipates the vesicular pH gradient and increases the vesicular membrane potential that VGLUT relies on for packaging glutamate [69]. And while we did not observe a reciprocal reduction in cholinergic transmission in the VGLUT1 cKO, this may be due to high variability in detection of cholinergic currents, or due to the presence of VGLUT2.

Nicotine consumption is shaped by a balance of its rewarding and aversive actions, thus our understanding of the circuit mechanisms by which nicotine aversion is encoded is crucial for developing effective therapeutics for nicotine addiction. Our findings demonstrate a role for glutamate signals from MHb cholinergic projections to IPN in opposing nicotine self-administration and suggests that potentiating nicotine’s effect on this circuit could be a useful target for nicotine cessation therapies. Future work may also focus on dissecting the relative roles of glutamate, ACh or other co-transmitters in this circuit on other aspects of nicotine behavior or in mediating responses to other substances of abuse.

## AUTHOR CONTRIBUTIONS

TSH conceived of the project. EAS, YC, VZ, YL, TS, WSC, WW, KDH, CDF and TSH contributed to data acquisition and analysis. EAS, WSC, CDF and TSH wrote the manuscript with input by all authors.

## REFERENCES

1 Wang TW, Gentzke AS, Creamer MR, Cullen KA, Holder-Hayes E, Sawdey MD, et al. Tobacco Product Use and Associated Factors Among Middle and High School Students — United States, 2019. MMWR Surveillance Summaries. 2019;68(12):1–22.

2 Kasza KA, Ambrose BK, Conway KP, Borek N, Taylor K, Goniewicz ML, et al. Tobacco-Product Use by Adults and Youths in the United States in 2013 and 2014. New England Journal of Medicine. 2017;376(4):342–53.

3 Cooper SY, Henderson BJ. The Impact of Electronic Nicotine Delivery System (ENDS) Flavors on Nicotinic Acetylcholine Receptors and Nicotine Addiction-Related Behaviors. Molecules. 2020;25(18):4223.

4 Dani JA, Heinemann S. Molecular and Cellular Aspects of Nicotine Abuse. Neuron. 1996;16(5):905–08.

5 Mansvelder HD, McGehee DS. Cellular and synaptic mechanisms of nicotine addiction. Journal of Neurobiology. 2002;53(4):606–17.

6 Wittenberg RE, Wolfman SL, De Biasi M, Dani JA. Nicotinic acetylcholine receptors and nicotine addiction: A brief introduction. Neuropharmacology. 2020;177:108256.

7 Akers AT, Cooper SY, Baumgard ZJ, Casinelli GP, Avelar AJ, Henderson BJ. Upregulation of nAChRs and Changes in Excitability on VTA Dopamine and GABA Neurons Correlates to Changes in Nicotine-Reward-Related Behavior. eNeuro. 2020;7(5):ENEURO.0189-20.

8 Calabresi P, Lacey MG, North RA. Nicotinic excitation of rat ventral tegmental neurones in vitro studied by intracellular recording. British Journal of Pharmacology. 1989;98(1):135–40.

9 Grieder TE, Besson M, Maal-Bared G, Pons S, Maskos U, Van Der Kooy D. β2* nAChRs on VTA dopamine and GABA neurons separately mediate nicotine aversion and reward. Proceedings of the National Academy of Sciences. 2019;116(51):25968–73.

10 Liu L, Zhao-Shea R, Mcintosh JM, Gardner PD, Tapper AR. Nicotine Persistently Activates Ventral Tegmental Area Dopaminergic Neurons via Nicotinic Acetylcholine Receptors Containing α4 and α6 Subunits. Molecular Pharmacology. 2012;81(4):541–48.

11 Mansvelder HD, McGehee DS. Long-Term Potentiation of Excitatory Inputs to Brain Reward Areas by Nicotine. Neuron. 2000;27(2):349–57.

12 Peng C, Engle SE, Yan Y, Weera MM, Berry JN, Arvin MC, et al. Altered nicotine reward-associated behavior following α4 nAChR subunit deletion in ventral midbrain. PLOS ONE. 2017;12(7):e0182142.

13 Pons S, Fattore L, Cossu G, Tolu S, Porcu E, Mcintosh JM, et al. Crucial Role of α4 and α6 Nicotinic Acetylcholine Receptor Subunits from Ventral Tegmental Area in Systemic Nicotine Self-Administration. Journal of Neuroscience. 2008;28(47):12318–27.

14 Pontieri FE, Tanda G, Orzi F, Chiara GD. Effects of nicotine on the nucleus accumbens and similarity to those of addictive drugs. Nature. 1996;382(6588):255–57.

15 Antolin-Fontes B, Ables JL, Görlich A, Ibañez-Tallon I. The habenulo-interpeduncular pathway in nicotine aversion and withdrawal. Neuropharmacology. 2015;96(Pt B):213-22.

16 Antolin-Fontes B, Li K, Ables JL, Riad MH, Görlich A, Williams M, et al. The habenular G-protein–coupled receptor 151 regulates synaptic plasticity and nicotine intake. Proceedings of the National Academy of Sciences of the United States of America. 2020;117(10):5502–09.

17 Fowler CD, Kenny PJ. Nicotine aversion: Neurobiological mechanisms and relevance to tobacco dependence vulnerability. Neuropharmacology. 2014;76:533–44.

18 Fowler CD, Kenny PJ. Intravenous nicotine self-administration and cue-induced reinstatement in mice: Effects of nicotine dose, rate of drug infusion and prior instrumental training. Neuropharmacology. 2011;61(4):687–98.

19 Fowler CD, Lu Q, Johnson PM, Marks MJ, Kenny PJ. Habenular α5 nicotinic receptor subunit signalling controls nicotine intake. Nature. 2011;471(7340):597–601.

20 Frahm S, Antolin-Fontes B, Görlich A, Zander JF, Ahnert-Hilger G, Ibañez-Tallon I. An essential role of acetylcholine- glutamate synergy at habenular synapses in nicotine dependence. eLife. 2015;4:e11396.

21 Frahm S, Ślimak MA, Ferrarese L, Santos-Torres J, Antolin-Fontes B, Auer S, et al. Aversion to Nicotine Is Regulated by the Balanced Activity of β4 and α5 Nicotinic Receptor Subunits in the Medial Habenula. Neuron. 2011;70(3):522–35.

22 Salas R, Sturm R, Boulter J, De Biasi M. Nicotinic receptors in the habenulo- interpeduncular system are necessary for nicotine withdrawal in mice. Journal of Neuroscience. 2009;29(10):3014–18.

23 Tuesta LM, Chen Z, Duncan A, Fowler CD, Ishikawa M, Lee BR, et al. GLP-1 acts on habenular avoidance circuits to control nicotine intake. Nature Neuroscience. 2017;20(5):708–16.

24 Elayouby KS, Ishikawa M, Dukes AJ, Smith ACW, Lu Q, Fowler CD, et al. α3* Nicotinic Acetylcholine Receptors in the Habenula-Interpeduncular Nucleus Circuit Regulate Nicotine Intake. The Journal of Neuroscience. 2021;41(8):1779–87.

25 McGehee DS, Heath MJS, Gelber S, Devay P, Role LW. Nicotine enhancement of fast excitatory synaptic transmission in CNS by presynaptic receptors. Science. 1995;269(5231):1692–96.

26 Perry DC, Xiao Y, Nguyen HN, Musachio JL, Dávila-Garcí MI, Kellar KJ. Measuring nicotinic receptors with characteristics of α4β2, α3β2 and α3β4 subtypes in rat tissues by autoradiography. Journal of Neurochemistry. 2002;82(3):468–81.

27 Salas R, Pieri F, De Biasi M. Decreased signs of nicotine withdrawal in mice null for the β4 nicotinic acetylcholine receptor subunit. Journal of Neuroscience. 2004;24(45):10035–39.

28 Shih PY, Engle SE, Oh G, Deshpande P, Puskar NL, Lester HA, et al. Differential Expression and Function of Nicotinic Acetylcholine Receptors in Subdivisions of Medial Habenula. Journal of Neuroscience. 2014;34(29):9789–802.

29 Dineley-Miller K, Patrick J. Gene transcripts for the nicotinic acetylcholine receptor subunit, beta4, are distributed in multiple areas of the rat central nervous system. Molecular Brain Research. 1992;16(3-4):339-44.

30 Marks M, Pauly J, Gross S, Deneris E, Hermans-Borgmeyer I, Heinemann S, et al. Nicotine binding and nicotinic receptor subunit RNA after chronic nicotine treatment. The Journal of Neuroscience. 1992;12(7):2765–84.

31 Wolfman SL, Gill DF, Bogdanic F, Long K, Al-Hasani R, McCall JG, et al. Nicotine aversion is mediated by GABAergic interpeduncular nucleus inputs to laterodorsal tegmentum. Nature Communications. 2018;9(1):2710.

32 Contestabile A, Villani L, Fasolo A, Franzoni MF, Gribaudo L, Øktedalen O, et al. Topography of cholinergic and substance P pathways in the habenulo-interpeduncular system of the rat. An immunocytochemical and microchemical approach. Neuroscience. 1987;21(1):253–70.

33 Herkenham M, Nauta WJH. Efferent connections of the habenular nuclei in the rat. Journal of Comparative Neurology. 1979;187(1):19–47.

34 Harrington L, Viñals X, Herrera-Solís A, Flores A, Morel C, Tolu S, et al. Role of β4* Nicotinic Acetylcholine Receptors in the Habenulo-Interpeduncular Pathway in Nicotine Reinforcement in Mice. Neuropsychopharmacology. 2016;41(7):1790–802.

35 Aizawa H, Kobayashi M, Tanaka S, Fukai T, Okamoto H. Molecular characterization of the subnuclei in rat habenula. The Journal of Comparative Neurology. 2012;520(18):4051–66.

36 Ren J, Qin C, Hu F, Tan J, Qiu L, Zhao S, et al. Habenula “Cholinergic” Neurons Corelease Glutamate and Acetylcholine and Activate Postsynaptic Neurons via Distinct Transmission Modes. Neuron. 2011;69(3).

37 Faget L, Zell V, Souter E, McPherson A, Ressler R, Gutierrez-Reed N, et al. Opponent control of behavioral reinforcement by inhibitory and excitatory projections from the ventral pallidum. Nature Communications. 2018;9(1).

38 Chen E, Lallai V, Sherafat Y, Grimes NP, Pushkin AN, Fowler J, et al. Altered Baseline and Nicotine-Mediated Behavioral and Cholinergic Profiles in ChAT-Cre Mouse Lines. The Journal of Neuroscience. 2018;38(9):2177–88.

39 Zell V, Steinkellner T, Hollon NG, Warlow SM, Souter E, Faget L, et al. VTA Glutamate Neuron Activity Drives Positive Reinforcement Absent Dopamine Co-release. Neuron. 2020;107(5):864–73.

40 Madisen L, Zwingman TA, Sunkin SM, Oh SW, Zariwala HA, Gu H, et al. A robust and high-throughput Cre reporting and characterization system for the whole mouse brain. Nature Neuroscience. 2010;13(1):133–40.

41 Görlich A, Antolin-Fontes B, Ables JL, Frahm S, Ślimak MA, Dougherty JD, et al. Reexposure to nicotine during withdrawal increases the pacemaking activity of cholinergic habenular neurons. Proceedings of the National Academy of Sciences of the United States of America. 2013;110(42):17077–82.

42 Oh JD, Woolf NJ, Roghani A, Edwards RH, Butcher LL. Cholinergic neurons in the rat central nervous system demonstrated by in situ hybridization of choline acetyltransferase mRNA. Neuroscience. 1992;47(4):807–22.

43 Trifonov S, Houtani T, Hamada S, Kase M, Maruyama M, Sugimoto T. In situ hybridization study of the distribution of choline acetyltransferase mRNA and its splice variants in the mouse brain and spinal cord. Neuroscience. 2009;159(1):344–57.

44 Fremeau RT, Troyer MD, Pahner I, Nygaard GO, Tran CH, Reimer RJ, et al. The expression of vesicular glutamate transporters defines two classes of excitatory synapse. Neuron. 2001;31(2):247–60.

45 Barroso-Chinea P, Castle M, Aymerich MS, Pérez-Manso M, Erro E, Tuñon T, et al. Expression of the mRNAs encoding for the vesicular glutamate transporters 1 and 2 in the rat thalamus. Journal of Comparative Neurology. 2007;501(5):703–15.

46 Varoqui H, Schäfer MKH, Zhu H, Weihe E, Erickson JD. Identification of the differentiation-associated Na+/PI transporter as a novel vesicular glutamate transporter expressed in a distinct set of glutamatergic synapses. Journal of Neuroscience. 2002;22(1):142–55.

47 Vong L, Ye C, Yang Z, Choi B, Chua S, Lowell BB. Leptin Action on GABAergic Neurons Prevents Obesity and Reduces Inhibitory Tone to POMC Neurons. Neuron. 2011;71(1):142–54.

48 Sartor CE, Lessov-Schlaggar CN, Scherrer JF, Bucholz KK, Madden PA, Pergadia ML, et al. Initial response to cigarettes predicts rate of progression to regular smoking: findings from an offspring-of-twins design. Addict Behav. 2010;35(8):771–8.

49 Dong Y, Zhang T, Li W, Doyon WM, Dani JA. Route of nicotine administration influences in vivo dopamine neuron activity: habituation, needle injection, and cannula infusion. J Mol Neurosci. 2010;40(1-2):164–71.

50 Corrigall WA, Coen KM. Nicotine maintains robust self-administration in rats on a limited- access schedule. Psychopharmacology (Berl). 1989;99(4):473–8.

51 Henningfield JE, Goldberg SR. Nicotine as a reinforcer in human subjects and laboratory animals. Pharmacol Biochem Behav. 1983;19(6):989–92.

52 Zevin S, Gourlay SG, Benowitz NL. Clinical pharmacology of nicotine. Clin Dermatol. 1998;16(5):557–64.

53 File SE, Cheeta S, Kenny PJ. Neurobiological mechanisms by which nicotine mediates different types of anxiety. Eur J Pharmacol. 2000;393(1-3):231-6.

54 Spealman RD. Maintenance of behavior by postponement of scheduled injections of nicotine in squirrel monkeys. J Pharmacol Exp Ther. 1983;227(1):154–9.

55 Risinger FO, Oakes RA. Nicotine-induced conditioned place preference and conditioned place aversion in mice. Pharmacol Biochem Behav. 1995;51(2-3):457–61.

56 Girod R, Barazangi N, Mcgehee D, Role LW. Facilitation of glutamatergic neurotransmission by presynaptic nicotinic acetylcholine receptors. Neuropharmacology. 2000;39(13):2715–25.

57 Hashikawa Y, Hashikawa K, Rossi MA, Basiri ML, Liu Y, Johnston NL, et al. Transcriptional and Spatial Resolution of Cell Types in the Mammalian Habenula. Neuron. 2020;106(5):743-58.e5.

58 Wallace ML, Huang KW, Hochbaum D, Hyun M, Radeljic G, Sabatini BL. Anatomical and single-cell transcriptional profiling of the murine habenular complex. Elife. 2020;9.

59 Hsu YW, Wang SD, Wang S, Morton G, Zariwala HA, de la Iglesia HO, et al. Role of the dorsal medial habenula in the regulation of voluntary activity, motor function, hedonic state, and primary reinforcement. J Neurosci. 2014;34(34):11366–84.

60 Hsu YW, Morton G, Guy EG, Wang SD, Turner EE. Dorsal Medial Habenula Regulation of Mood-Related Behaviors and Primary Reinforcement by Tachykinin-Expressing Habenula Neurons. eNeuro. 2016;3(3).

61 Otsu Y, Darcq E, Pietrajtis K, Matyas F, Schwartz E, Bessaih T, et al. Control of aversion by glycine-gated GluN1/GluN3A NMDA receptors in the adult medial habenula. Science. 2019;366(6462):250–54.

62 Pang X, Liu L, Ngolab J, Zhao-Shea R, McIntosh JM, Gardner PD, et al. Habenula cholinergic neurons regulate anxiety during nicotine withdrawal via nicotinic acetylcholine receptors. Neuropharmacology. 2016;107:294–304.

63 Soria-Gomez E, Busquets-Garcia A, Hu F, Mehidi A, Cannich A, Roux L, et al. Habenular CB1 Receptors Control the Expression of Aversive Memories. Neuron. 2015;88(2):306–13.

64 Yamaguchi R, Nicholson Perry K, Hines M. Pain, pain anxiety and emotional and behavioural problems in children with cerebral palsy. Disabil Rehabil. 2014;36(2):125–30.

65 Zhang J, Tan L, Ren Y, Liang J, Lin R, Feng Q, et al. Presynaptic Excitation via GABAB Receptors in Habenula Cholinergic Neurons Regulates Fear Memory Expression. Cell. 2016;166(3):716–28.

66 Zhao-Shea R, Liu L, Pang X, Gardner PD, Tapper AR. Activation of GABAergic neurons in the interpeduncular nucleus triggers physical nicotine withdrawal symptoms. Current Biology. 2013;23(23):2327–35.

67 Zhao-Shea R, DeGroot SR, Liu L, Vallaster M, Pang X, Su Q, et al. Increased CRF signalling in a ventral tegmental area-interpeduncular nucleus-medial habenula circuit induces anxiety during nicotine withdrawal. Nat Commun. 2015;6:6770.

68 Qin C, Luo M. Neurochemical phenotypes of the afferent and efferent projections of the mouse medial habenula. Neuroscience. 2009;161(3):827–37.

69 Hnasko TS, Edwards RH. Neurotransmitter corelease: mechanism and physiological role. Annu Rev Physiol. 2012;74:225–43.

